# A phosphoramidate modification of FUDR, NUC-3373, causes DNA damage and DAMPs release from colorectal cancer cells, potentiating lymphocyte-induced cell death

**DOI:** 10.1101/2024.10.09.617410

**Authors:** Oliver J. Read, Jennifer Bré, Ying Zhang, Peter Mullen, Mustafa Elshani, David J. Harrison

**Affiliations:** School of Medicine, University of St Andrews, North Haugh, St Andrews, Fife, UK. KY16 9TF; NuCana plc, 3, Lochside Way, Edinburgh, UK EH12 9DT; Present address: Jacqui Woods Cancer Centre, University of Dundee, DD2 1GZ

**Keywords:** Colorectal cancer, DAMPs, coculture, pro-drug, immunotherapy

## Abstract

Colorectal cancer (CRC) is one of the leading causes of cancer-related mortality worldwide with 5-FU still the primary chemotherapeutic of choice. With the increasing use of immunotherapies, much research is focused on the ability to make tumours more immunogenic, thereby rationalising combination with immunotherapy. Here we investigate whether NUC-3373, a phosphoramidate transformation of FUDR, enhances immunogenicity in CRC cell lines and facilitates lymphocyte mediated cell death *in vitro*. At sub IC_50_ doses NUC-3373 upregulates damage associate molecular patterns (DAMPs) in both HCT116 and SW480 cells and increases surface expression of MHCII and PD-L1. Pre-treatment with NUC-3373 and subsequent coculture with NK-92 MI natural killer cells caused an increase in LAMP1 expression (degranulation), production of IFN-γ, and NK-mediated cytotoxicity compared to vehicle controls. Cocultures with patient-derived PBMCs with heterologous CRC cells pre-treated with NUC-3373 demonstrated increased cell death compared to both vehicle controls and monocultures of CRC cells exposed to NUC-3373. Lastly, the PD-1 immune checkpoint inhibitor nivolumab showed synergistic activity when HCT116 cells were pre-treated with NUC-3373. To conclude, we show that NUC-3373 can modulate immune signaling and may therefore facilitate immune mediated tumour cell death.

## 1 Introduction

Colorectal cancer (CRC) remains one of the largest challenges to oncology with the third highest incidence and second highest mortality rate globally (10% and 9.4%, respectively)^1^. While chemotherapy remains the backbone of therapy for CRC^2,3^, in the last decade immunotherapy has stepped into the forefront of cancer treatment. Several immunotherapies have emerged that employ different approaches to either kick-start or prolong anti-tumour immunity and encourage immunogenic cell death (ICD). These range from adoptive cell transfer, including genetically modified autologous cells such as T-cells with chimeric antigen receptors (CAR), anti-cancer vaccines, and immune checkpoint inhibitors^4^, *e.g.,* antibodies that block programmed cell death protein 1 (PD-1), programmed cell death ligand 1 (PD-L1), or cytotoxic T lymphocyte antigen 4 (CTLA4).

Currently four monoclonal antibodies targeting immune checkpoints are approved by the FDA in CRC for treatment of tumours with deficient mismatch repair (dMMR) and/or high microsatellite instability (MSI)^5–9, 63^. These are nivolumab (anti-PD-1), pembrolizumab (anti-PD-1), atezolizumab (anti-PD-L1), and ipilimumab (anti-CTLA4). The rationale behind the use of such immunotherapies in dMMR/MSI tumours is based on the propensity of cancer cells to upregulate immune checkpoint molecules, tumour associated antigens (TAAs), and major histocompatibility complex (MHC) II molecules due to high mutational burden^2,9^. However, this only accounts for 3-6% of advanced stage CRC, with the majority being microsatellite stable (MSS) and/or mismatch repair proficient (MMR-p)^2,10,11^.

Immunotherapies present attractive candidates for combination with traditional anti-cancer approaches such as radiotherapy and chemotherapy agents, especially those that are known to increase immunogenicity of tumour cells. Cytotoxicity caused by such treatments can trigger the presentation and/or release of damage-associated molecular patterns (DAMPs) from stressed and dying cells which can be detected by circulating dendritic cells (DCs), causing differentiation into antigen presenting cells (APCs) that signal to other immune cells in the tumour microenvironment to trigger anti-tumour immune response or amplify an ongoing response^12,13^. Although several DAMPs have been described, the most immunogenic signals are the cell surface exposure of calreticulin (CRT) and heat-shock proteins (Hsp) alongside the release of high-mobility-group box 1 protein (HMGB1) and adenosine triphosphate (ATP)^14^. Whilst most chemotherapies provoke release of DAMPs, this is usually achieved using high doses of drug for very short time periods *in vitro*^15,16^.

5-fluorouracil (5-FU) remains one of the most widely prescribed anti-cancer drugs and is used to treat common cancers including colorectal, gastric, pancreatic and head and neck. 5-FU and other fluoropyrimidines are pro-drugs that require conversion to the active metabolite 5-fluorodeoxyuridine-monophosphate (FUDR-MP or FdUMP), which binds and inhibits thymidylate synthase (TS)^17^. TS is required to convert deoxyuridine monophosphate (dUMP) to deoxythymidine monophosphate (dTMP)^17,18^. Blocking TS results in an imbalance in the ratio dUMP and dTMP, with subsequent depletion of deoxythymidine triphosphate (dTTP) and disruption of DNA synthesis and repair, ultimately leading to apoptosis. However, several key limitations impacting the bioavailability and activation of 5-FU have been associated with sub-optimal responses to treatment^17,19^. The majority (>85%) of administered 5-FU is degraded by the enzyme dihydropyrimidine dehydrogenase (DPD) in the liver, generating alpha-fluoro-beta-alanine (FBAL), a catabolite associated with off-target toxicities such as hand-foot syndrome. The 5-FU that is taken up by cancer cells is dependent on expression of thymidine phosphorylase and thymidine kinase for conversion to FUDR and phosphorylation to FUDR-MP. Thus, alteration in the levels of these enzymes limits the anti-cancer activity of 5-FU. Furthermore, misincorporation of the metabolite fluorodeoxyuridine triphosphate (FUTP) in RNA is partly responsible for efficacy but also causes myelosuppression and gastrointestinal toxicity^20–22^.

NUC-3373, a novel anti-cancer ProTide, is a phosphoramidate transformation of 5-fluorodeoxyuridine (FUDR) comprised of FUDR and a phosphoramidate moiety^23–25^. The phosphoramidate protects the molecule from DPD-mediated degradation, conferring reduced exposure to toxic catabolites and associated toxicities, as well as significantly improving the plasma half-life compared to 5-FU (6-10 hours versus 8-14 minutes)^26^. As NUC-3373 contains the pre-phosphorylated anti-cancer metabolite, there is no need for enzymatic conversion to FUDR and subsequent phosphorylation. Once inside the cancer cell, the protective groups are cleaved with release of significantly higher levels of the active anti-TS metabolite, FUDR-MP, compared to 5-FU^25^. Additionally, we have shown that in addition to inhibition of thymidylate synthase, NUC-3373 exerts cytotoxicity through a DNA-targeted mode of action through misincorporation of FUDR into DNA, resulting in DNA damage and cell cycle arrest, and negligible misincorporation of fluorouridine (FUR) into RNA compared to 5-FU^25^.

This study investigated the pro-immunogenic effects of NUC-3373 in CRC cell lines by investigating cellular events independent of its cytotoxic mode of action, in particular ER stress and DAMPs release, importantly using doses less than IC_50_. Using *in vitro* cocultures of CRC cells with either a natural killer (NK) cell line or volunteer-derived peripheral blood mononuclear cells (PBMCs) we show that NUC-3373 can modulate the pro-inflammatory signaling and increase tumour cell death which is complemented after addition of a PD-1 checkpoint inhibitor.

## 2 Materials and Methods

### 2.1 PBMC isolation from whole blood

PBMCs were isolated from whole blood samples from healthy volunteers approved by the University of St Andrews School of Medicine Ethics Committee (approval code: MD15716). Whole blood was drawn into vacutainer tubes containing EDTA and diluted 1:1 with sterile PBS. Blood solution was then layered onto 10 ml of Histopaque 1077 (Sigma Aldrich) and centrifuged at 800 x g for 30 minutes (no brake). Buffy coat was transferred to a separate tube and resuspended in PBS before centrifuging at 300 x g for 10 minutes. Two additional wash steps with PBS were performed before resuspending cells in freeze media (10% DMSO in FBS) and storing at -80°C until required.

### 2.2 Cell culture and reagents

HCT116 (microsatellite instable) and SW480 (microsatellite stable) cell lines were cultured in Dulbecco’s Modified Eagle Medium (DMEM – Gibco) with 10% (v/v) fetal bovine serum and 1% (v/v) Penicillin/Streptomycin. Cells were incubated at 37°C with 5% CO_2_ and media was replaced every 3-4 days. NK-92 MI cells were cultured in alpha-minimal essential media (MEM - Gibco) supplemented with 12.5% horse serum (v/v), 12.5% fetal bovine serum (v/v), 1% Penicillin/Streptomycin, 200 μM myo-inositol (Sigma Aldrich), 100 μM β-mercaptoethanol (Sigma Aldrich), and 20 μM folic acid (Sigma Aldrich). Cells were incubated at 37°C with 5% CO_2_. Cell lines routinely tested negative for Mycoplasma using the Minerva Biolabs ‘Venor GeM One Step’ PCR kit.

NUC-3373 was supplied as a powder by NuCana plc and was dissolved in DMSO (Sigma Aldrich) to achieve a stock concentration of 40 mM and stored at -20°C. Stock NUC-3373 was diluted to a 10 mM working concentration before use. LPS and PHA (Sigma Aldrich) were dissolved in DMSO to give concentrations of 0.5 and 2.5 mg/ml respectively. PMA and ionomycin (Sigma Aldrich) were dissolved in DMSO to achieve stock concentrations of 500 μg/ml and 5 mM respectively. Oxaliplatin (Sigma Aldrich) was dissolved in deionised water to a stock concentration of 10 mM. 5-FU (Sigma Aldrich) was dissolved in DMSO to a stock concentration of 10 mM and stored at -20°C. Thymidine powder (Sigma Aldrich) was dissolved in dH_2_O to achieve a final concentration of 50 μg/ml. A working solution of thymidine was made by diluting stock in DMEM to make an 8 μg/ml solution.

### 2.3 Immunofluorescence microscopy

Cells were plated onto poly-d-lysine coated coverslips in 6-well plates and left to adhere for 48 hours before treatment. For intracellular staining of HMGB1 and γ-H2AX, cells were fixed with 4% paraformaldehyde and permeabilised with 0.1% Triton-X-100 in PBS. Aldehyde sites were blocked with 0.1 M glycine, following which coverslips were washed with PBS then blocked in 10% goat serum in PBS for 10 minutes at RT. Coverslips were incubated with primary antibody (Table 1) for 1 hour at RT. Coverslips were washed with PBS then incubated with secondary antibody for 45 minutes in the dark at RT. Coverslips were then washed with PBS-Tween before incubation with Hoechst 33342 (Invitrogen) in PBS (1:10,000 dilution) for 10 minutes at RT. Cells were then washed with PBS-Tween and rinsed with dH_2_O. Coverslips were dried and mounted onto slides using Prolong Gold antifade mountant (Invitrogen).

For fluorescent labelling of intracellular ATP with quinacrine, a stock 10 mM quinacrine solution was diluted to 10 µM in cell culture media. Quinacrine-containing media was then aliquoted onto live cells grown on coverslips and incubated for 20 minutes at RT in the dark. Cells were washed with PBS before nuclear staining with Hoechst in PBS for 10 minutes at RT in the dark. Coverslips were washed twice with PBS and mounted on slides with Prolong Gold antifade mountant. Images were taken with a Leica CTR5500 epifluorescence microscope (40x magnification). Image analysis was performed with Image J.

### 2.4 Flow Cytometry

HCT116 and SW480 cells were seeded in 6-well plates at 15,000 and 37,500 cells/well respectively and were allowed to attach and grow for 48 hours. Cells were then treated with NUC-3373, vehicle control (DMSO – dilution dependent on maximal dose of NUC-3373), or oxaliplatin depending on specific assay and cell line and incubated for 24 hours. CRC cells were then either harvested immediately or drug containing media was removed to allow for PBMC coculture. For cocultures 1 x 10^6^ PBMCs were added to CRC-containing wells and incubated until harvesting 24 and 48 hours later. Antibodies used for flow cytometry are described in Table 1. In instances where cells needed to be fixed, samples were incubated for 15 minutes in 4% formaldehyde in PBS at room temperature then moved to 4°C until required. Acquisition of samples was performed using a CytoFlex (Beckman Coulter) and analysis performed using CytExpert (ver 2.4) or FlowJo (ver 10.8.1). Gating strategies for each experiment are listed in supplementary.

### 2.5 NK degranulation and cytokine accumulation assays to determine activation

CRC cells were plated and treated as described previously. Drug-containing media was removed after 24 hours exposure and NK-92 MI cells resuspended in fresh DMEM were added at a density of 2 × 10^5^ cells per well and incubated for 4 hours before harvesting the suspension cells (NK cells) for flow cytometry. For the positive control, co-cultured NK-92 MI cells were treated with PMA/ionomycin at 50 ng/ml & 0.5 μM respectively for 4 hours. Negative control constituted untreated NK-92 MI cells alone. Full details of antibodies are presented in Table 1: in summary, for assessment of degranulation, 5 μl of anti-LAMP1 was added to each well at the start of coculture. For assessment of IFN-γ production, GolgiStop (BD Biosciences) was added to DMEM containing NK cells as per the manufacturer’s instructions. After the 4-hour coculture period suspension cells were harvested and resuspended and washed in 0.5% BSA in PBS and stained with anti-CD56. Samples being studied for intracellular IFN-γ were fixed in 4% formaldehyde/PBS for 15 minutes at room temperature then permeabilised with 0.3% Triton-X100 in PBS.

### 2.6 CRC viability assays in NK or PBMC cocultures

HCT116 and SW480 cells were dispensed onto 96-well plates at 500 and 1500 cells/well respectively and placed in an incubator for 48 hours to allow for adherence and growth. Media was aspirated and fresh media was added containing 1, 5, 10 μM NUC-3373 for HCT116 cells and 10, 25, 40 μM for SW480 cells and incubated for a further 24 hours. DMSO at 0.1 and 0.4% was used as a vehicle control for HCT116 and SW480 cells respectively. Before initiation of coculture, PBMCs were prepared at 67,000 cells/well in DMEM. Half of the PBMCs were treated with 10 μg/ml of nivolumab. Drug-containing media was removed from the 96-well plate and replaced with either plain DMEM, DMEM containing PBMCs, or DMEM containing PBMCs and nivolumab. Plates were then placed in an incubator for up to 72 hours with separate plates being allocated for 24, 48, and 72 hours of coculture. Confluence was assessed every 24 hours starting at the time of addition of drug using a Celigo scanner (Nexelcom Biosciences). At the end of the coculture period a Sulforhodamine B (SRB) assay was performed to determine cell viability as previously described^27^. Plates were scanned using a BP800 Microplate Reader (540 nm absorbance filter).

For NK coculture assays CRC cells were treated with either vehicle control (DMSO), 5/10 μM NUC-3373 for HCT116 or 10/25 μM NUC-3373 for SW480 cells. After 24 hours drug-containing media was removed and replaced with either fresh media or media containing 1.3 x 10^4^ NK-92 MI cells for coculture. Separate plates were fixed at specific time points before continuing with SRB protocol.

### 2.7 Nanopore RNA-seq

HCT116 and SW480 cells were seeded in 10cm dishes at 88,600 and 265,800 cells/dish respectively and were allowed to attach and grow for 48 hours. Cells were then treated with NUC-3373(15uM for HCT116 and 35uM for SW480), 5-FU (15uM for HCT116 and 35uM for SW480) or vehicle control (DMSO – dilution dependent on dose of NUC-3373) and incubated for 24 hours. CRC cells were then either harvested immediately or drug containing media was changed to normal culture media to allow cell grow for another 24h or 48h before harvesting. Then cells were lysed and RNA was isolated and purified using RNeasy mini kit (Qiagen) according to the manufacturer’s instructions. cDNA libraries were performed from 500 ng total RNA according to the Oxford Nanopore Technologies PCR-cDNA Barcoding Kit (SQK-PCB111.24) protocol with a 16 cycles PCR. Nanopore libraries were sequenced using a PromethION 2 Solo with R10.4.1 flow cells (Oxford Nanopore Technologies Ltd). We utilised the nanopore P2 sequencer with PromethION flow cells, following the manufacturer’s protocol with the cDNA-based PCB114.24 ONT transcriptomics kits. Base calling was performed using Dorado v0.7.0 on 4 A100 GPUs. The base called reads were mapped to the Gencode v41 transcriptome using minimap2 v2.28, and transcript abundance was quantified using Salmon v1.10 with the long-read --ont parameter. Differential gene and transcript expression analysis was conducted using DESeq2 and DRIMSeq, respectively.

### 2.8 Statistical analysis

Analysis of flow cytometry data, including gating, was performed using either FlowJo (ver 10.6.1) or CytExpert (ver 2.4) software. Raw values for generation of figures were exported to Microsoft Excel and analysed in GraphPad (ver 8.4). Additional analysis is detailed in figure legends.

## 3 Results

### 3.1 NUC-3373 promotes DAMPs release at sub-IC50 concentration

Flow cytometric analysis revealed a 45% increase of calreticulin (CRT) at the cell surface of HTC116 cells after 24 hours of exposure to NUC-3373 at less than IC50 levels, compared to untreated cells (Fig 1A.) and a >40% increase for SW480 cells at an equimolar dose (Fig 1B.). Treatment with NUC-3373 also resulted in loss of nuclear HMGB1 after 24 hours, which became more pronounced at 48 hours (Fig 1C.). Change in HMGB1 localisation in SW480 cells at equivalent dose was less, likely due to SW480 being a more resistant cell line to NUC-3373 meaning fewer cells would be undergoing pre-/apoptotic HMGB1 release (Fig 1D.). Finally, intracellular ATP was labelled with quinacrine for fluorescence microscopy as an indirect qualitative assay for ATP release (Fig. 1E.). After 24 hours of treatment with NUC-3373, the pattern of ATP staining shifted from diffuse to more punctate, characteristic of a shift to increased vesicular ATP, consistent with literature describing subsequent release from cells^28^. Treatment of SW480 cells with 10 μM NUC-3373 also produced similar punctate staining of ATP in cells (Fig 1F.). Additionally, NUC-3373 caused an increase in surface expression of Hsp70 (Fig S1.), a non-classical DAMP, in both cell lines

**Figure 1.**
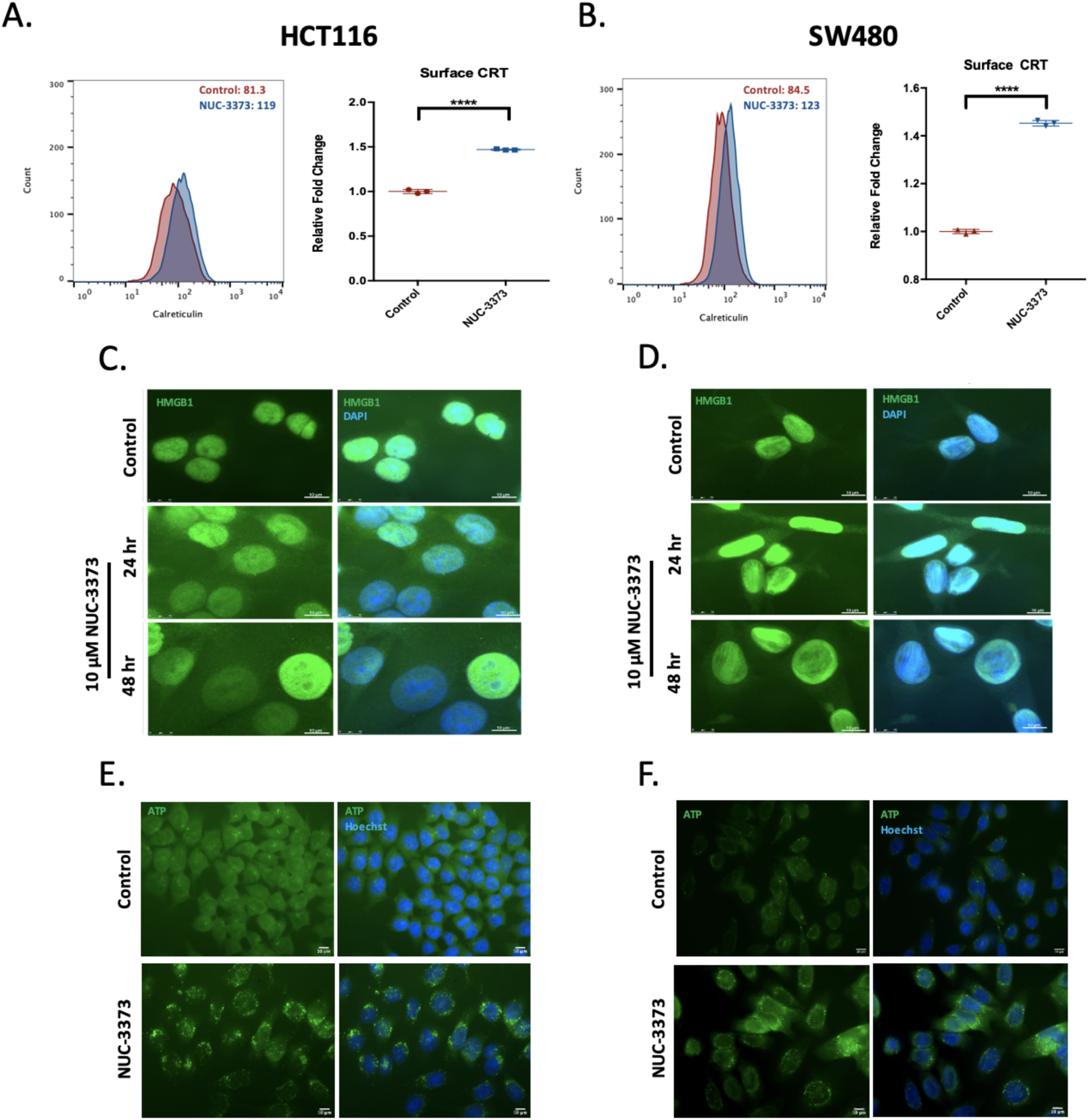
NUC-3373 causes DAMPs release in HCT116 cells and SW480 cells. **A. &B.** Flow cytometry analysis of cell-surface CRT in HCT116 and SW480 cells respectively treated with NUC-3373 for 24h (blue) compared to DMSO control (red) with accompanying fold changes based on median fluorescent intensity (MFI). Graph displays mean and standard deviation of MFI fold change across three biological replicates. MFI for respective replicate displayed on representative histograms for HCT116 and SW480 cells. **C. & D.** Immunofluorescence images staining for HMGB1 (green) and counterstained with Hoechst (blue) at 24h and 48h post-treatment with NUC-3373 or vehicle control. **E. & F.** Imaging of intracellular ATP through use of quinacrine fluorescent labelling. Hoechst (blue) was used as a nuclear counterstain. Mean and standard deviation based on the relative fold change of three replicates. Significant difference between control and treated cells was determined by Student’s T-test (**** p < 0.0001).

### 3.2 Differential expression of immune-related genes between NUC-3373 and 5-FU treated cells

Comprehensive RNA Seq analysis conducted over a time course of 72 hours is ongoing, but in HCT-116 and SW480 colorectal cancer cell lines similar patterns were observed for NUC-3373, generating three clusters that were distinct from changes induced by 5-FU (Figure 2). These changes were upregulation of HLA DR, beta 2-microglobulin and PDL-1; downregulation of histone related genes, H1, 2 and 4; and upregulation of thymidylate synthase and thymidine kinase, the latter being important in the thymidine salvage pathway.

**Figure 2.**
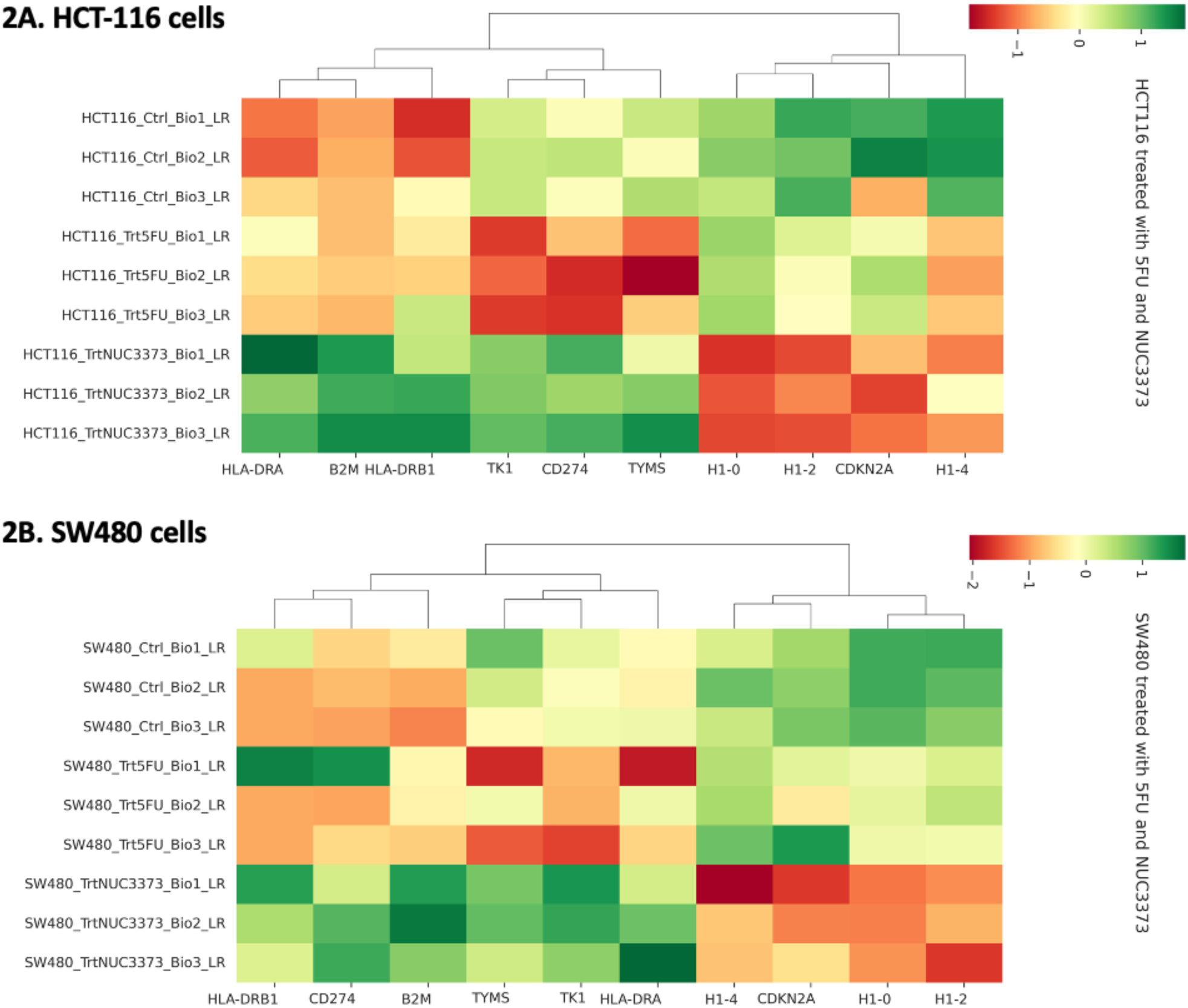
Summary of mRNA changes in three colorectal cancer cell lines 72 hours after treatment with near-IC50 doses of NUC-3373 and 5-FU. Figures show heatmaps for three biological replicates for each of control, 5-FU and NUC-3373 treated HCT116 cells (2A) and SW480 (2B). These changes are shown at 72 hours but a time course was conducted (results not shown). Differences in gene expression between 5-FU and NUC-3373 are illustrated particularly affecting HLA DR, beta 2-microglobulin, histone proteins H1,2 and 4, and thymidylate synthase and thymidine kinase. Some of these changes have subsequently been confirmed in biopsies from patients taken before and after treatment with NUC-3373.

### 3.3 NUC-3373 pre-treatment potentiates the activation of NK cells

To assess whether NUC-3373-associated release and exposure of DAMPs would potentiate tumour cell death via innate immunity, CRC cells were cocultured with a NK cell line (NK-92 MI^29–31)^. Activation of NK cells can be characterised both *in vitro* (and *in vivo)* through assessment of degranulation (release of cytolytic vesicles), determined through quantification of surface-exposed LAMP1 (CD107a)^32–34^. NK activity can further be characterised *in vitro* through changes in cytokine production, such as accumulation of interferon gamma (IFN-γ) after being treated with a Golgi transport inhibitor to prevent cytokine release. HCT116 cells were exposed to 10 μM NUC-3373 for 24 hours. After washing out the drug, CRC cells were cocultured with NK cells for 4 hours. NK cells were then assessed for degranulation and intracellular IFN-γ by flow cytometry. NK cells cocultured with NUC-3373-treated CRC cells displayed a 125% increase in LAMP1 surface expression and a 96% increase in intracellular IFN-γ relative to DMSO-treated controls (determined using median fluorescent intensity – Fig 3A. & C.). In both assays, NUC-3373 outperformed oxaliplatin, a known inducer of ICD^35^. In SW480 cells, there was an increase in both LAMP1 and IFN-γ levels upon treatment with higher doses of NUC-3373 (40 μM). When cocultured with SW480 cells exposed to the higher dose of NUC-3373, LAMP1 surface expression of NK cells increased by 57% (Fig 3B.) compared to DMSO control, meanwhile intracellular IFN-γ levels only increased by 8% (Fig 3D.).

**Figure 3.**
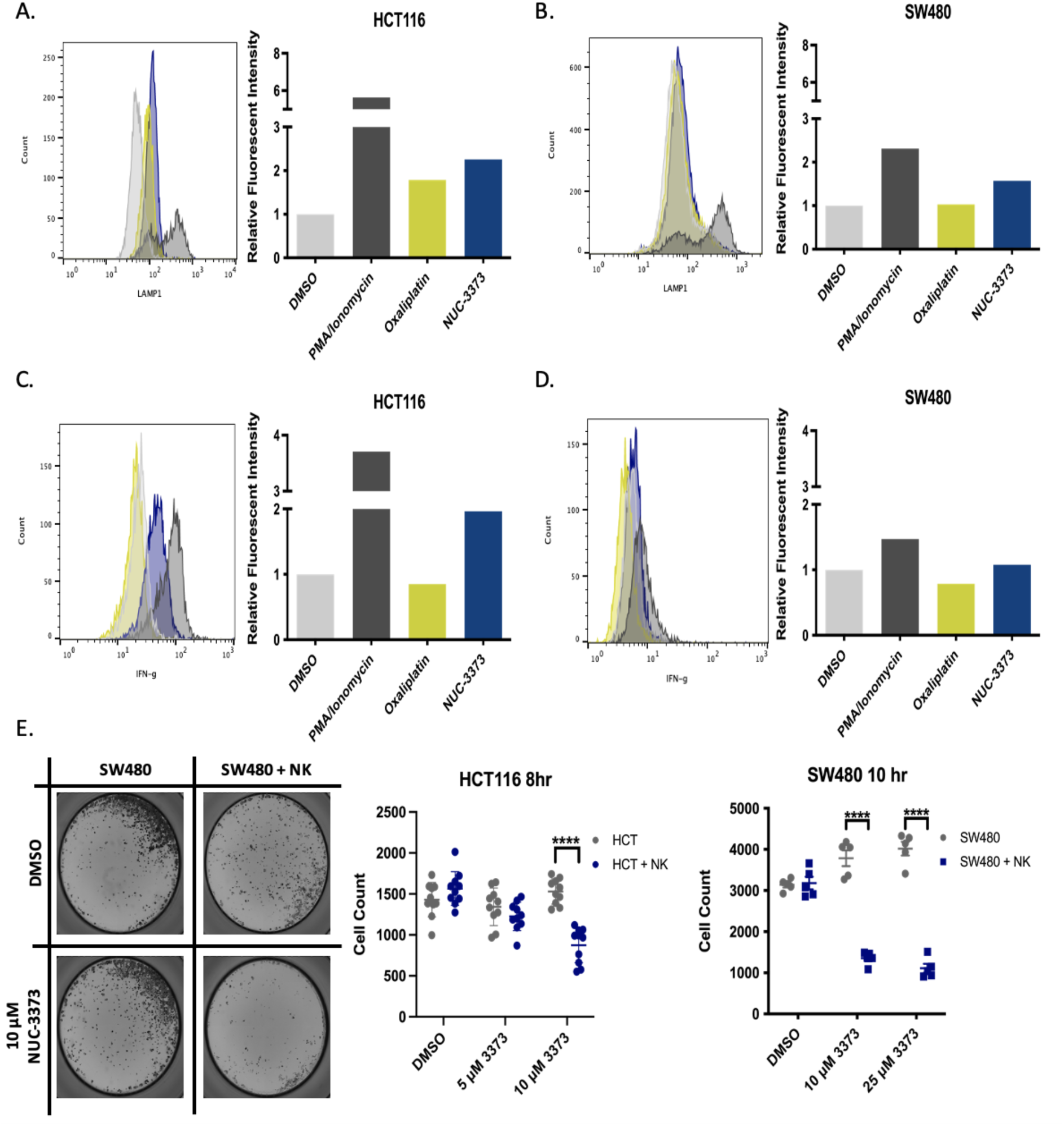
NUC-3373 pre-treatment of CRC cells causes NK cell activation: Flow cytometric analysis of **A. & B.** surface LAMP1 expression and **C. & D.** intracellular IFN-γ for NK-92 MI cells cocultured with HCT116 (left) or SW480 (right) that had been pre-treated with either DMSO control, 10 μM oxaliplatin (yellow), or 10 μM NUC-3373 (blue). PMA/ionomycin (dark grey) was used as a positive control. Figures show representative plots, additional replicates are shown in Fig S5. **E.** CRC cells in 96-well plates were exposed to either DMSO or NUC-3373 for 24 hours and cultured for up to an additional 10 hours in the presence or absence of NK-92 MI cells. Following removal of NK cells and SRB staining, plates were imaged (left – representative wells for SW480) and analysed by automated cytometry analysis to determine cell count (middle and right graphs). Plots show mean with standard deviation of all wells of two biological replicates. Difference between means determined by Student’s t-test (**** p < 0.0001).

With observed increases in NK activation markers, coculture cytotoxicity assays were performed to assess if DAMP release caused by NUC-3373 treatment would increase NK-mediated cytotoxicity. CRC cells were exposed to different concentrations of NUC-3373 in 96-well plates for 24 hours after which drug was removed. Plates were scanned and cell counts in the presence or absence of NK cells was determined by automated cytometry analysis after SRB staining. For both cell lines, pre-treatment with NUC-3373 caused a significant decrease in cell counts in coculture conditions (Fig 3E.) indicating an increase in ICD due to NUC-3373-mediated DAMPs release.

### 3.4 NUC-3373 pre-treatment causes upregulation of pro-immune signals and also increases PD-L1 in CRC cells

After confirming that NUC-3373 was able to stimulate NK cells and promote an innate anti-tumour response *in vitro* we assessed whether DAMPs release from CRC cells due to NUC-3373 had a similar effect on the adaptive immune system using a similar coculture model. HCT116 cells were pre-treated with NUC-3373 before coculture with volunteer-derived PBMCs for 24-48 hours. Analysis of stimulatory cytokines by qPCR revealed that NUC-3373 pre-treatment resulted in a dramatic increase in mRNA of IL-2, TNF-α, and IFN-γ at 24 hours (Fig S3A), especially in conditions where PBMCs were stimulated with anti-CD3/28 where it appears that NUC-3373 amplified the effect of the stimulatory antibodies by further increasing gene expression of said cytokines. mRNA expression of PD-L1 was also measured in co-cultures and was found to be upregulated in samples pre-treated with NUC-3373 compared to DMSO controls (Fig S3A.). This increase in PD-L1 was also confirmed at the protein level by measuring surface PD-L1 of both HCT116 and PBMC cells by flow cytometry (Fig S3B.)

As PD-L1 is a checkpoint inhibitor molecule that is often upregulated by cancer cells to promote immune evasion^36^, additional experiments were performed to confirm if the observed increase is due to increased immune signaling (i.e., cytokine secretion from immune cells) or if it is a more direct effect of NUC-3373. In addition to upregulation of PD-L1 through exogenous immune cytokine signaling such as IFN-γ, PD-L1 is also upregulated in response to DNA damage via induction of ataxia telangiectasia and Rad3-related protein (ATR) and subsequent phosphorylation of checkpoint kinase 1 (Chk1)^37,38^. Previous work established that, compared to 5-FU, NUC-3373 treated CRC cells have higher levels of misincorporated FUDR in DNA which results in increase in DNA-damage markers yH2AX and p-Chk1 and cell cycle arrest^25^. A head-to-head comparison of 5-FU and NUC-3373 reveals that in both HCT116 and SW480 monocultures, NUC-3373 causes an increase in cell-surface PD-L1 but 5-FU does not at the concentrations (Fig 4A. & B). We then replicated the results seen in the previous study, however using IF, whereupon treatment with NUC-3373 caused an increase in intracellular levels of y-H2AX with only a small increase in signal being observed for equimolar concentrations of 5-FU in both cell lines (Fig 4C. & D). Based on this evidence from CRC monocultures, it is likely that the increase in PD-L1 expression is largely due to DNA damage caused by NUC-3373.

**Figure 4.**
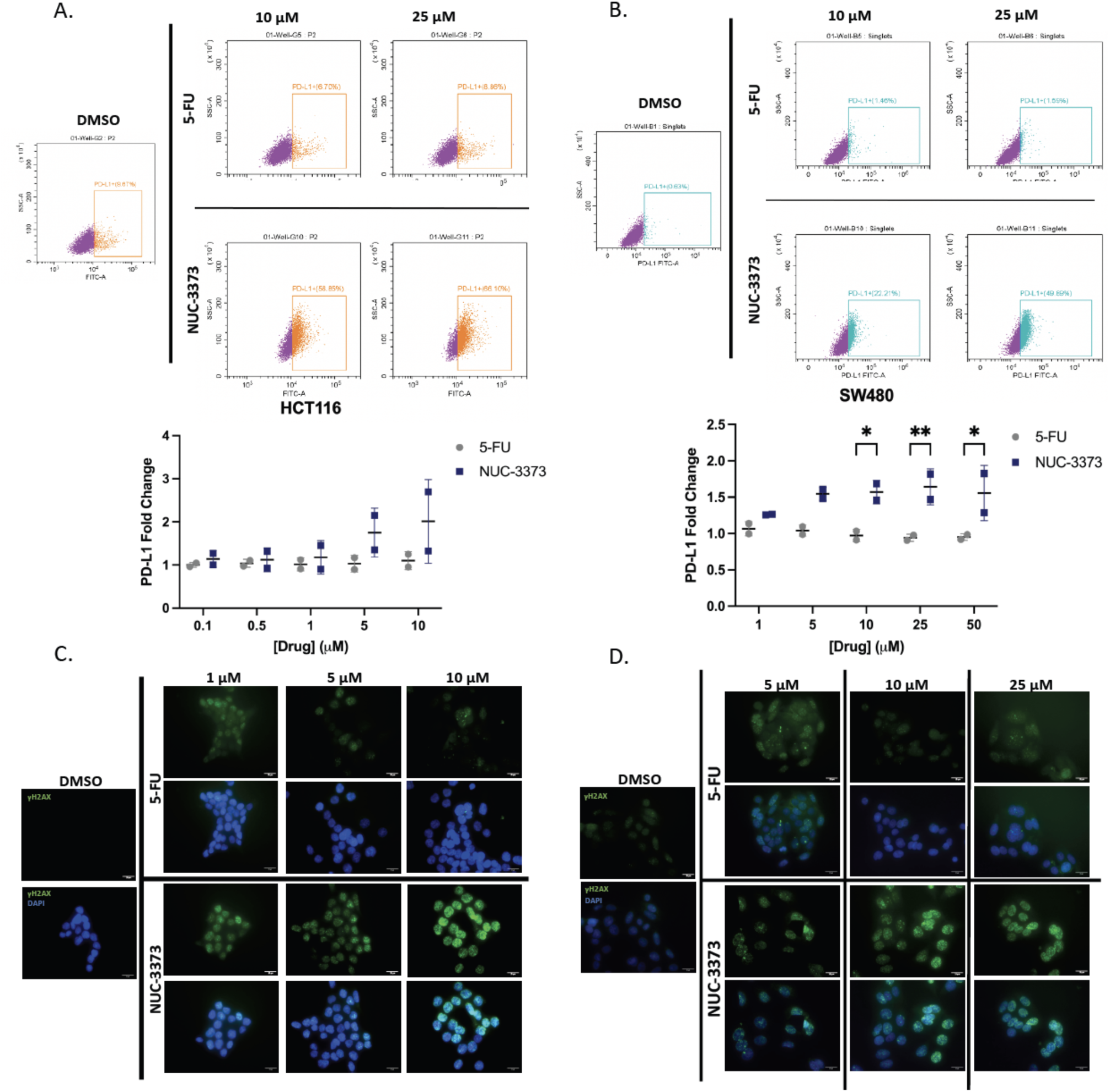
NUC-3373 increases surface expression of PD-L1 in CRC cells and DNA damage: HCT116 and SW480 cells were treated with increasing doses of 5-FU and NUC-3373 (0.1 – 10 μM and 1 – 50 μM for each cell line respectively) or vehicle control (DMSO) for 24 hours before harvesting for flow cytometry. **A. & B.** representative flow cytometry plots (top) and corresponding summary graph for each test condition in HCT116 and SW480 respectively following staining of cell-surface PD-L1. Gates indicate cells positive for surface PD-L1 staining. Graphs display data from individual biological replicates (n=2) with mean and standard deviation. Statistical significance was determined by two-way ANOVA (* p<0.05, ** p<0.01). **C. & D.** Immunofluorescent staining for γ-H2AX in HCT116 and SW480 cells respectively treated with either vehicle control (DMSO), 5-FU, or NUC-3373 at different concentrations.

**Figure 5.**
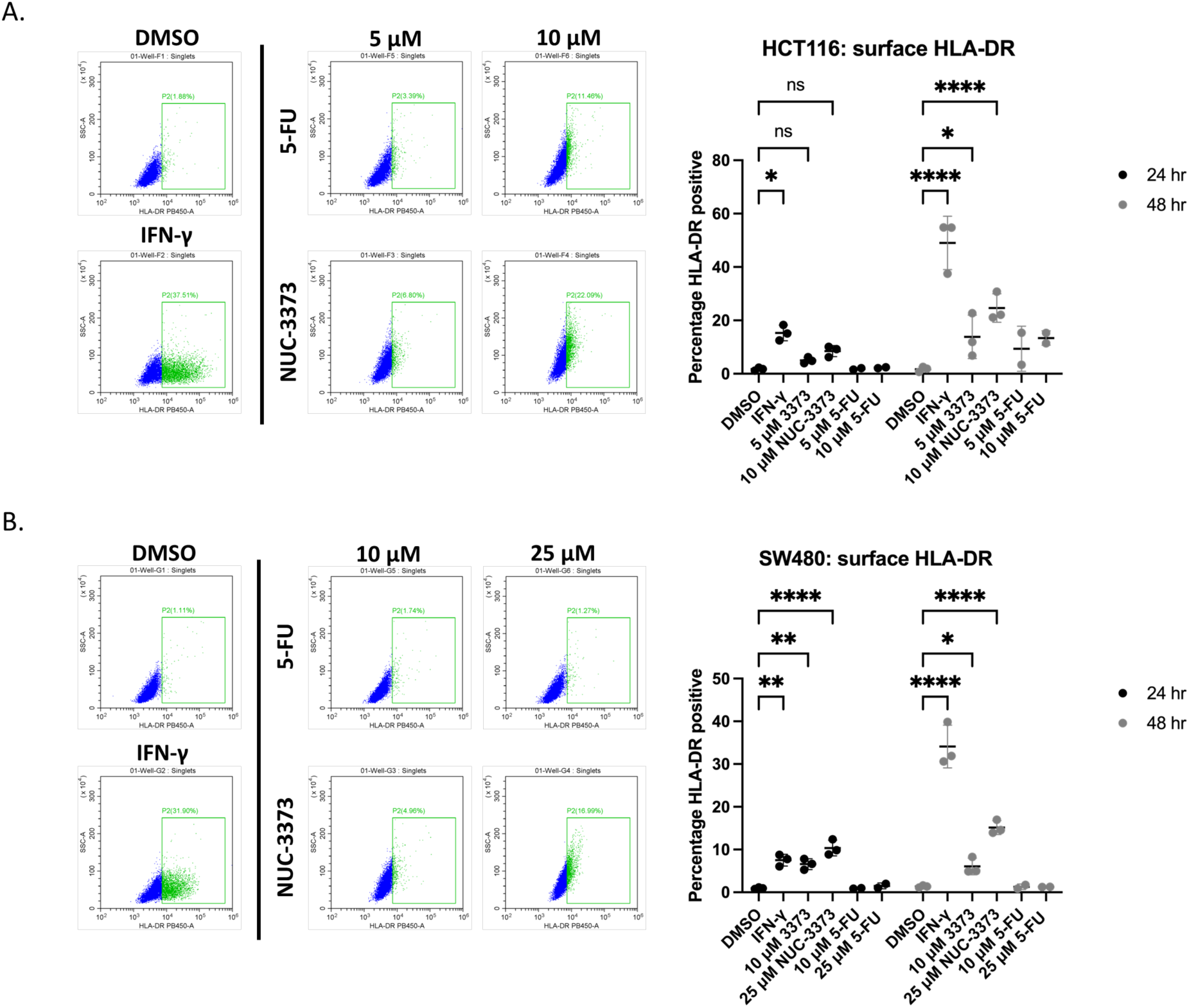
NUC-3373 causes increase in surface HLA-DR expression in CRC. HCT116 (**A.**) and SW480 (**B.**) cells were treated with vehicle control (DMSO), IFN-γ (20 ng/ml), or different doses of NUC-3373/5-FU and analysed by flow cytometry for surface expression of HLA-DR at 24 and 48 hours. Representative flow plots on left showing HLA-DR against side scatter for each condition at 48 hours, right graphs display data for 3 biological replicates for NUC-3373 and 2 biological replicates for 5-FU at both time points including mean and standard deviation. Statistical significance between DMSO and treatment conditions determined by 2-way ANOVA (* = p < 0.5, ** = p < 0.01, **** = p < 0.0001).

DNA damage has also been implicated in the upregulation of MHC class I and II (MHCI/II)^39,40^. Although studies have mainly focused on induction and upregulation of MHCI, Lhuillier and colleagues have shown that radiotherapy also increases neoantigen presentation from MHCII in a triple negative breast cancer model^39^. As DNA damage-induced MHCI involves ATR signaling, similar to PD-L1 we hypothesised that NUC-3373 could also upregulate MHCII expression. Flow cytometric analysis of HLA-DR, an inducible type of MHCII the expression of which may be a favorable prognostic marker in CRC^41^, on HCT116 and SW480 monocultures revealed that NUC-3373 caused a significant upregulation of surface HLA-DR expression by 48 hours of treatment in a concentration-dependent manner, with SW480 cells showing significance after 24 hrs relative to DMSO control. Similar to what was observed with PD-L1 levels, treatment with equimolar concentrations of 5-FU resulted in only a modest increase in surface HLA-DR after 48 hrs treatment in HCT116 cells.

### 3.5 NUC-3373 primes PBMCs for immune-mediated cytotoxicity of CRC cells

To address these seemingly paradoxical results of pro-immune (i.e., DAMP release, more potent cytokine induction, increased NK activation, and NK-mediated cytotoxicity) and anti-immune (i.e., increased PD-L1 mRNA and surface protein) readouts, a series of further coculture experiments were performed with PBMCs to determine whether NUC-3373 can promote lymphocyte mediated tumour cell death. Nivolumab was added in some conditions at the time of coculture to test the hypothesis that NUC-3373 in combination with a PD-1/PD-L1 checkpoint inhibitor would be synergistic in a coculture environment. Cell viability of CRC cell lines was assessed in a variety of different treatment conditions in the presence or absence of PBMCs so that drug-specific effects on cell viability can be compared with any additional cytotoxicity caused by PBMCs or PD-1 inhibitor.

For HCT116 cells, presence of PBMCs resulted in decreased viability of cells treated with vehicle control at both 48 and 72 hours with no significant difference when nivolumab was added (Fig 6A. & B. – grey bars). Treatment with 5 and 10 μM NUC-3373 caused a drop in cell viability at both time points compared to vehicle control (Fig 6B. – blue and purple bars). Coculture with PBMCs in these conditions resulted in statistically significant drop in viability for 5 μM NUC-3373 at both time points with no difference observed for 10 μM. However, whilst addition of nivolumab had no effect for 5 μM at 48 hours compared to the coculture condition, checkpoint blockade yielded a decrease in viability at 72 hours compared to other 5 μM-treated conditions. Furthermore, addition of nivolumab for 10 μM pre-treatment resulted in significant drop in cell viability at both time points compared to HCT116 monoculture conditions.

**Figure 6.**
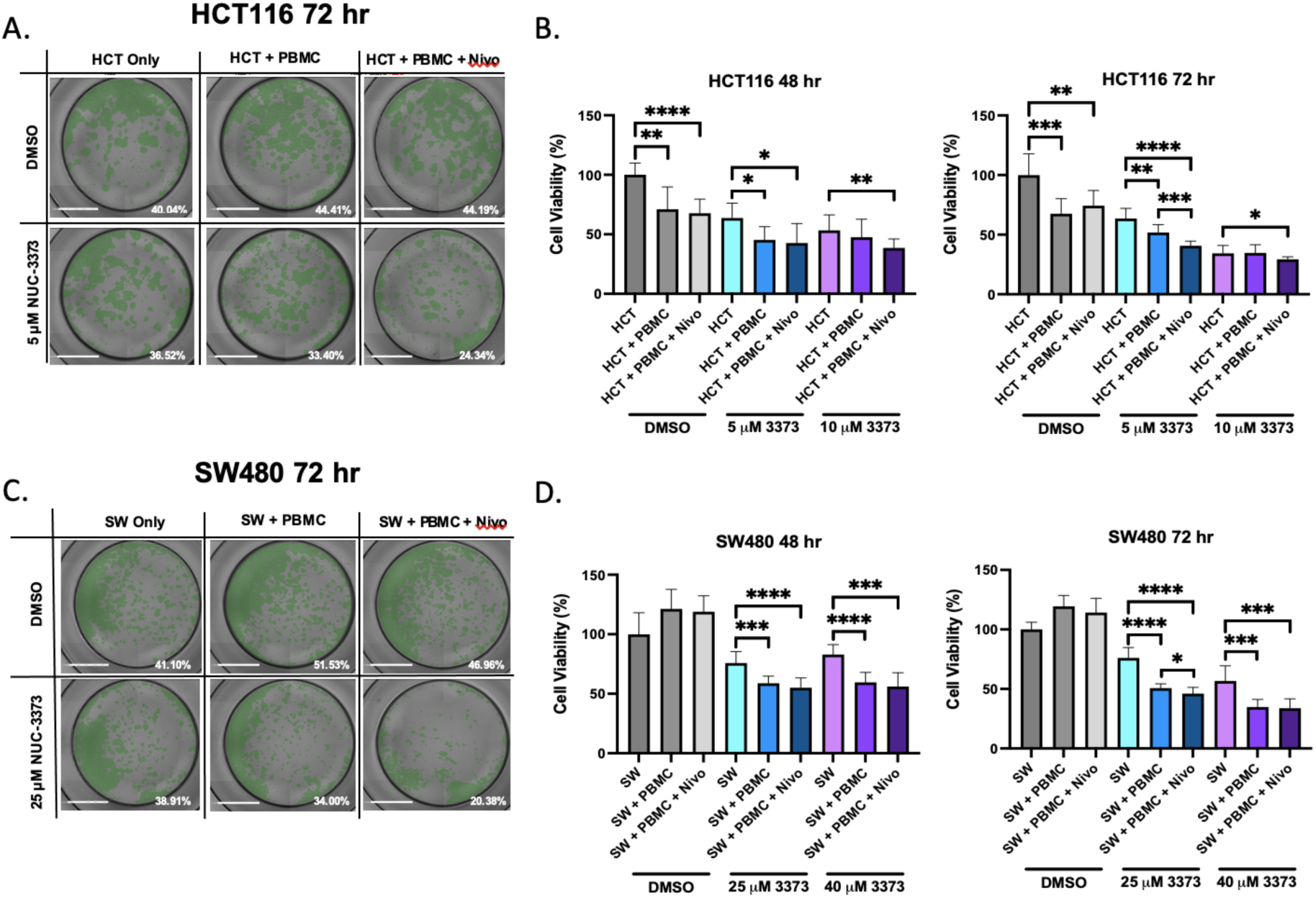
NUC-3373 potentiates PBMC-mediated cytotoxicity of CRC cells in coculture: HCT116 or SW480 were seeded onto 96-well plates and cocultured with patient-derived PBMCs after 24 hours exposure to NUC-3373 or vehicle control (DMSO). Some wells were additionally treated with 10 µg/ml nivolumab (Nivo) at the start of co-culture. Plates were imaged and analysed by both automated cytometry analysis and SRB assay. **A.** representative images of HCT116 cells in wells after 72 hrs of coculture for vehicle control and 5 µM NUC-3373. Well confluence for respective well is displayed in the bottom right of each image. **B.** Cell viability of HCT116 cells at 48 hrs (left) and 72 hrs (right) of co-culture, determined by SRB. **C.** Representative images of SW480 cells after 72 hrs co-culture for vehicle control and 25 µM. **D.** Cell viability data for SW480 cells determined by SRB. SRB data was normalised to CRC cell line only with vehicle control test condition. Graphs and error display mean and standard deviation of 5 wells across 2 biological replicates. Statistical significance was determined by Mann-Whitney test (* p<0.05, ** p<0.01, *** p<0.001, **** p<0.0001).

Interestingly for SW480 cells, presence of PBMCs appeared to increase overall cell viability of CRC cells for the vehicle control (Fig 6D. – grey bars). Pre-treatment with 25 and 40 μM NUC-3373 caused a decrease in cell viability in conditions with coculture of PBMCs decreasing viability even further (Fig 6D. – blue and purple bars). Addition of nivolumab appeared to make no difference to cell viability except for a small significant difference for 25 μM pre-treatment at 72 hours coculture.

## 4 Discussion

In this study we show that NUC-3373-mediated inhibition of TS and leads to release of DAMPs in CRC cell lines. HLA DR and b2 microglobulin increased and we found potentiated activation and cytotoxic effects when coculturing NK cells with treated tumour cells. In CRC/PBMC cocultures, increased gene expression of pro-immune cytokines including IL-2, IFN-γ and TNF-α was observed in cases where CRC cells had been pre-treated with NUC-3373. NUC-3373 also caused an increase in PD-L1 mRNA expression and MHCII, raising the intriguing prospect that NUC-3373 may be a good combination partner for PD-(L)1 targeted checkpoint therapy. Additional experiments verified that pre-treatment of cells with NUC-3373 caused increased cytotoxicity in CRC cells cocultured with PBMCs compared to both vehicle control and monoculture conditions and that addition of the anti-PD1 checkpoint inhibitor nivolumab causes further decrease in cell viability.

In addition to NUC-3373’s ability to act as a cytotoxic through inhibition of TS and misincorporation of FUDR into DNA we hypothesised that NUC-3373 could also promote the release of DAMPs from CRC cells, an event that may favour immune mediated cell death^13^.^41^ Many cytotoxics cause varying degrees of DAMPs release, often detected after exposure to high doses *in vitro* within a very short window of time^16^. In this study, DAMPs release was observed in cells exposed to a sub-IC_50_ dose, at concentrations of drug that have been reported in patients^64^, supporting that this may also occur *in vivo*. Interaction of cancer cells with immune cells in the tumour microenvironment is a key factor in patient prognosis and the efficacy of anti-cancer therapies. Some chemotherapeutics are reported to have greater efficacy *in vivo* in immune-competent organisms as opposed to immune-compromised organisms or *in vitro*^13,42^. The cytotoxic activity of chemotherapeutics or radiotherapy can promote the release of pro-immunogenic DAMPs from dying cells, which is detected by circulating NK cells as well as dendritic cells that undergo maturation and can develop into antigen presenting cells (APCs)^13,43,44^. In both cell lines examined, increased cell surface CRT expression along with loss of nuclear HMGB1 after 24 hours suggests that NUC-3373 activates DAMP pathways in CRC cells. In addition, the shift in ATP staining from diffuse to more punctate following treatment with NUC-3373 is characteristic of ATP packaging in vesicles prior to exocytosis^16,45^. These observations indicate that NUC-3373 may have the ability to induce pro-immunogenic changes in the tumour microenvironment. Definitive evidence that is the case would require *in vivo* studies however, mice have high levels of esterases which rapidly degrade phosphoramidate moieties^46–48^, so no *in vivo* experiments have been conducted, but ongoing studies in patients’ biopsies before and after treatment have been undertaken (results not shown).

NK-92 is an immortalised natural killer cell line that has been used to model the acute response of innate immunosurveillance^29,49,50^. The variant of NK-92 used in this study was modified to produce its own interleukin-2 (IL-2) to allow self-replication rather than requiring IL-2 supplementation of culture media. Whilst NK cells do not respond to as many DAMPs as dendritic cells, they are receptive to HMGB1 as they express Toll-like receptor 4^51,50^ in addition to surface expression of Hsp70 on damaged cells^51–53^. In brief, following a 4-hour coculture incubation period, NK cells cultured with CRC cells that had been previously treated with NUC-3373 showed increased surface expression of LAMP1 and production of IFN-γ, which are classical markers of NK cell activation^32,33^. NK activation was associated with increased NK cytotoxic activity in both cell lines pre-treated with NUC-3373. It is important to note that as NUC-3373 was not present in media during the coculture period, any differences in NK activity would solely be due to NUC-3373-associated DAMPs being released within the timeframe.

During experiments involving PBMC/CRC cocultures there was an increase in both mRNA expression and cell surface protein of PD-L1, despite the observed increase in DAMP signalling and gene expression of pro-immune cytokines. This could be induced by either the presence of IFN-γ in the media released by PBMCs, as indeed IFN-γ is often used as a positive control for induction of PD-L1 in cell lines *in vitro*, and/or DNA damage. Activation of DNA damage response (DDR) mechanisms due to cell cycle arrest or strand breaks can increase expression of PD-L1 via ATR/ATM signaling resulting in phosphorylation of Chk-1, STAT3 activation and upregulation of interferon response protein^52–54^. Inhibition of thymidine synthesis and increased fluorodeoxyuridine incorporation into DNA following NUC-3373 treatment causes cell cycle arrest and upregulation of γH2AX and p-Chk1^55^, which we corroborated in this study. Indeed, studies have also shown this phenomenon with 5-FU in both p53 wild-type and mutant HCT116 cells in monocultures, however at significantly higher doses ^56^ than those used in this study where more clinically achievable doses were used. We also observed increase in MHCII molecule HLA-DR, a favorable prognostic marker in CRC^41,57^, when CRC cells were exposed to increasing concentrations of NUC-3373, which was more pronounced in HCT116 cells, which may be due to MMR status. Typically, HLA-DR is expressed by professional antigen-presenting cells (such as dendritic cells and monocytes) however their expression can be induced in tumour and epithelial cells when exposed to inflammation^41,58^. Although there is a significant degree of literature that attests to DNA damaging agents increasing the expression of MHCI, DNA damage from radiotherapy has also been shown to increase expression of MHCII^39^ which was the basis behind this line of investigation. Induction of MHCII has been shown to elicit prolonged anti-tumour immune response as it promotes activation of CD4+ cells^59^. Studies have shown the importance of co-stimulation of both CD8+ and CD4+ T-cells for both spontaneous and immunotherapy-triggered anti-tumour immune responses and for circumventing immune tolerance of cancer cells^39,59,60^.

Upregulation of PD-L1 in CRC cells could therefore be a response to both increased pro-immune microenvironment changes but in particular DNA damage due to drug treatment. This presented an opportunity to investigate the *in vitro* efficacy of a PD-1/PD-L1 axis inhibitor in PBMC/CRC cocultures as we hypothesised that addition of a checkpoint inhibitor would increase lymphocyte-related tumour cell death by overcoming any inhibitory effect of PD-L1 upregulation. Indeed, an increase in tumour cell death with nivolumab treatment was observed in HCT116 cells when comparing different conditions. By contrast the addition of nivolumab to SW480 cells had only a slightly significant effect at 72 hours with 25 μM NUC-3373 pre-treatment. Pre-treatment with NUC-3373 caused a significant drop in cell viability upon coculture with PBMCs relative to both vehicle controls and CRC monoculture controls, suggesting that NUC-3373 facilitates PBMCs ability to target the underlying CRC cells, irrespective of the cell lines mismatch repair and microsatellite stability background. This raises the intriguing possibility of conversion of immunogenically “cold” tumours to being amenable to immune checkpoint therapy, not just the 3-6% of patients with d-MMR/MSI. As we saw an increase in PD-L1 expression upon treatment with NUC-3373, it may initially seem that an anti-PD-L1 inhibitor such as durvalumab would have been more appropriate over nivolumab. However, whilst durvalumab has seen promise as a monotherapy in phase II clinical trials in MSI/dMMR CRC ^61^, in theory blocking the PD-1/PD-L1 axis at any point might elicit a similar effect.

Interestingly, addition of PBMCs to SW480 cells in DMSO conditions increased cell viability compared to monoculture, which contrasts with what was observed for HCT116 cells, where coculture with PBMCs was sufficient for decreasing cell viability. It was therefore noteworthy to see that pre-treatment with NUC-3373 caused a significant drop in cell viability compared to the monoculture control, especially considering that SW480 is an MMR-p/MSS cell line^62^. It is also encouraging to be able to conclude that addition of a PD-1 inhibitor can only have positive effects in terms of efficacy, with no apparent antagonism in any conditions involving NUC-3373.

We attempted to control for MHC incompatibility in our system by including vehicle control and monoculture conditions to establish cell death due to MHC mismatch, this is potentially why we observe significant cell death in HCT116 cells treated with DMSO when cocultured with PBMCs. Using primary tumour cells or tissue organoids with autologous PBMCs would eliminate MHC incompatibility as a confounder.

Establishment of molecular mechanisms and general pharmacology *in vitro* is a crucial step in the drug development process; however, it is also important to try and establish the downstream effects of the candidate compounds mode of action in a wider context than just the target cell. In addition to traditional *in vivo* studies using animal models to identify potential toxicities and immunological impact on host organisms, *in vitro* studies have begun to focus more on factors that can influence the tumour microenvironment. The data presented here demonstrate that NUC-3373 has the potential to modulate the tumour microenvironment to facilitate anti-tumour immune activity. It also sets a strong precedent for combination with checkpoint inhibitors and potentially other forms of immunotherapy. The finding that NUC-3373 in combination with nivolumab was able to potentiate tumour cell death after addition of lymphocytes, irrespective of the cell lines’ MSI status, indicates the potential for a wide clinical utility for NUC-3373 in the treatment of cancer.

## 6 Author Contributions

OJR and DJH conception and design of research. OJR, JB, PM, ME, and YZ performed experiments and analysed data. OJR., ME., PM and DJH. interpreted results. OJR. and ME drew figures. OJR drafted manuscript. All authors edited and revised the final version of the manuscript.

## Acknowledgments

We would like to thank Dr Danielle McLean for assistance in writing.

