## supplemental data for "A phosphoramidate modification of FUDR, NUC-3373, causes DNA damage and DAMPs release from colorectal cancer cells, potentiating lymphocyte-induced cell death"

### Supplementary Methods:

#### Gene expression of cytokines and PD-L1

HCT116/PBMC cocultures were set up in 6-well plates as previously described with the amount of PBMCs added after drug incubation being 1 x $10^6$ cells. RNA lysates were prepared at 24 and 48 hours coculture time points using a RNeasy Kit (Qiagen) as per manufacturer's instructions. Extracted RNA was then converted to cDNA using a QuantiTect Reverse Transcription Kit (Qiagen). Primers for qPCR included TNF-α: fwd- GCTGCACTTTGGAGTGATCG, rev- GCTGAGGGTTTGCTACAA; IFN-γ: fwd- CGTTTTGGGTTCTCTTGGCT, rev- TTTCTGTCACTCTCCTCTTTCC; IL-2: fwd- TTACATGCCCAACAAGGCCA, rev- TGGTTGCTGTCTCATCAGCAT; and PD-L1: fwd- AGGCCGAAGTCATCTGGACAAG, rev- TCCTCTCTCTTGGAATTGGTG (ThermoFisher) and QuantiTect Primer Assays for GAPDH (Qiagen) and β-Actin (Qiagen). Samples for qPCR were prepared using QuantiNova SYBR green PCR Kit (Qiagen) and run on a Rotor-Gene Q (Qiagen) and associated software. Ct data was exported to and analysed in Excel using the ΔΔCt method.

#### Antibodies

| Antibody/Target | Conjugate | Application | Supplier | Catalogue no. |
|---|---|---|---|---|
| β-Actin | UC | WB (1:2,000) | Cell Signaling Technology | 3700s |
| Donkey anti-rabbit | IRDye 800RD | WB (1:10,000) | Licor | 926-32213 |
| Donkey anti-mouse | IRDye 680RD | WB (1:10,000) | Licor | 926-68072 |
| HMGB1 | UC | IF | Abcam | 18256 |
| γ-H2AX | UC | IF | Cell Signalling Technology | 9718 |
| Hsp-70 | UC | FC (2.5 μg/ml) | Biolegend | 648002 |
| Goat anti-mouse | FITC | FC (1:10,000) | Abcam | 6785 |
| CRT | UC | FC (1:800) | Cell Signaling Technology | 12238 |
| Goat anti-rabbit | AlexaFluor 488 | FC (1:10,000) | Cell Signaling Technology | 4412 |
| CD17a (LAMP1) | AlexaFluor 488 | FC (5 μl/1 x 10e6 cells) | Biolegend | 328610 |
| CD56 | AlexaFluor 647 | FC (5 μl/1 x 10e6 cells) | Biolegend | 318314 |
| TIGIT | PE | FC (5 μl/1 x 10e6 cells) | Biolegend | 373703 |
| IFN-γ | AlexaFluor 488 | FC (5 μl/1 x 10e6 cells) | Biolegend | 505813 |

| | | | | |
|---|---|---|---|---|
| CD274 (PD-L1) | FITC | FC (5 µl/1 x 10e6 cells) | Biolegend | 374510 |
| CD3 | UC | Stim (1 µg/ml) | Biolegend | 300302 |
| CD28 | UC | Stim (1 µg/ml) | Biolegend | 302902 |
| Cytokeratin 7/8 | AlexaFluor 647 | FC (5 µl/1 x 10e6 cells) | BD Biosciences | 563614 |
| HLA-DR | Pacific Blue | FC (5 µl/1 x 10e6 cells) | Biolegend | 307633 |
| Nivolumab (anti-PD-1) | UC | AB (10 µg/ml) | Selleckchem | A2002 |

**Table 1:** List of antibodies and applications. Abbreviations: **UC** – unconjugated; **WB** - western blot; **IHC** – immunohistochemistry; **IF** – immunofluorescence; **FC** – flow cytometry; **AB** – antigen blocking.

### Supplementary Figures:

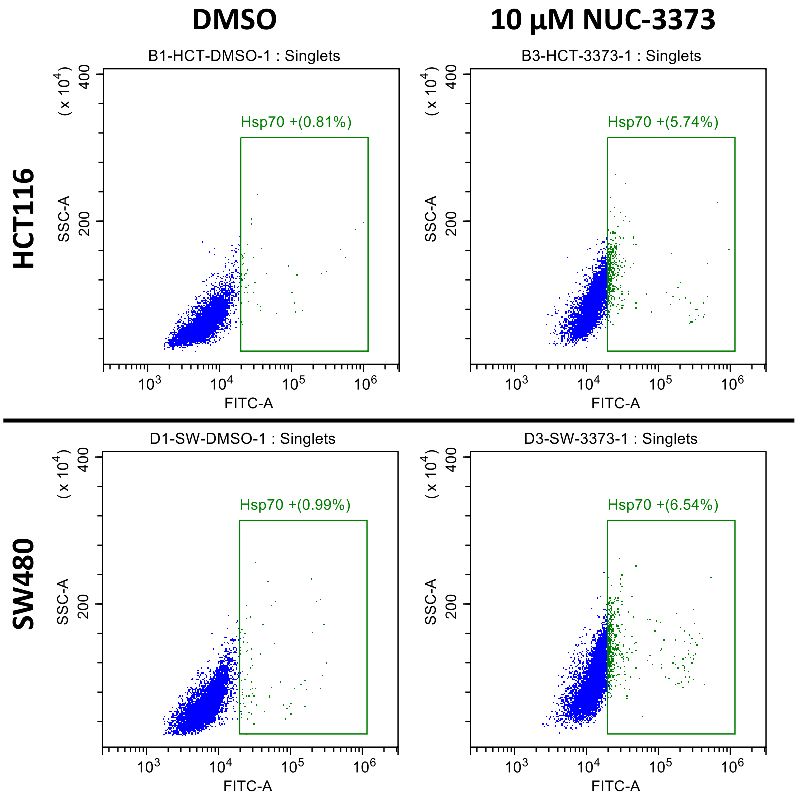


**Fig S1. NUC-3373 causes increased surface expression of Hsp-70 on CRC cells:** Flow cytometry data for Hsp-70 on HCT116 (top) and SW480 (bottom) cells after being exposed to vehicle control (DMSO – left) or 10 µM NUC-3373 (right) for 24 hours. Gates drawn based on unstained controls.

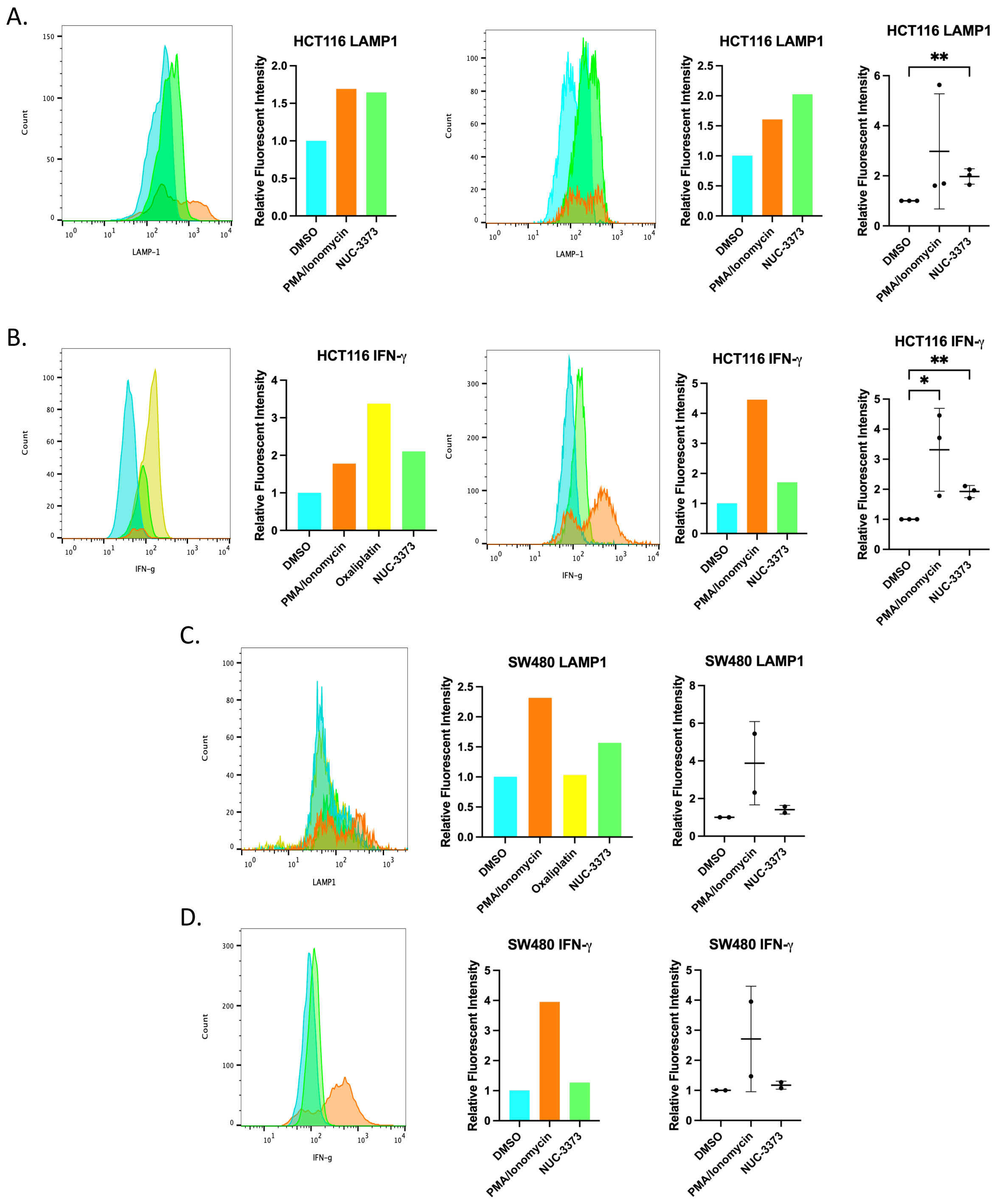


**Fig S2. Additional replicates for LAMP1 and IFN-γ levels for NK cells cocultured with CRC cells:** flow histograms, normalised fluorescent intensity for additional replicates and summary data for all reps for LAMP1 and IFN-γ in NK cells cocultured with pre-treated HCT116 (**A.** & **B.**) and SW480 (**C.** & **D.**) cells. Media fluorescent intensity was normalised to that of DMSO control within each experiment. Summary plots show individual datapoints, mean, and standard deviation. Difference between means was analysed by Student's t-test (* = p < 0.5, ** = p < 0.01).

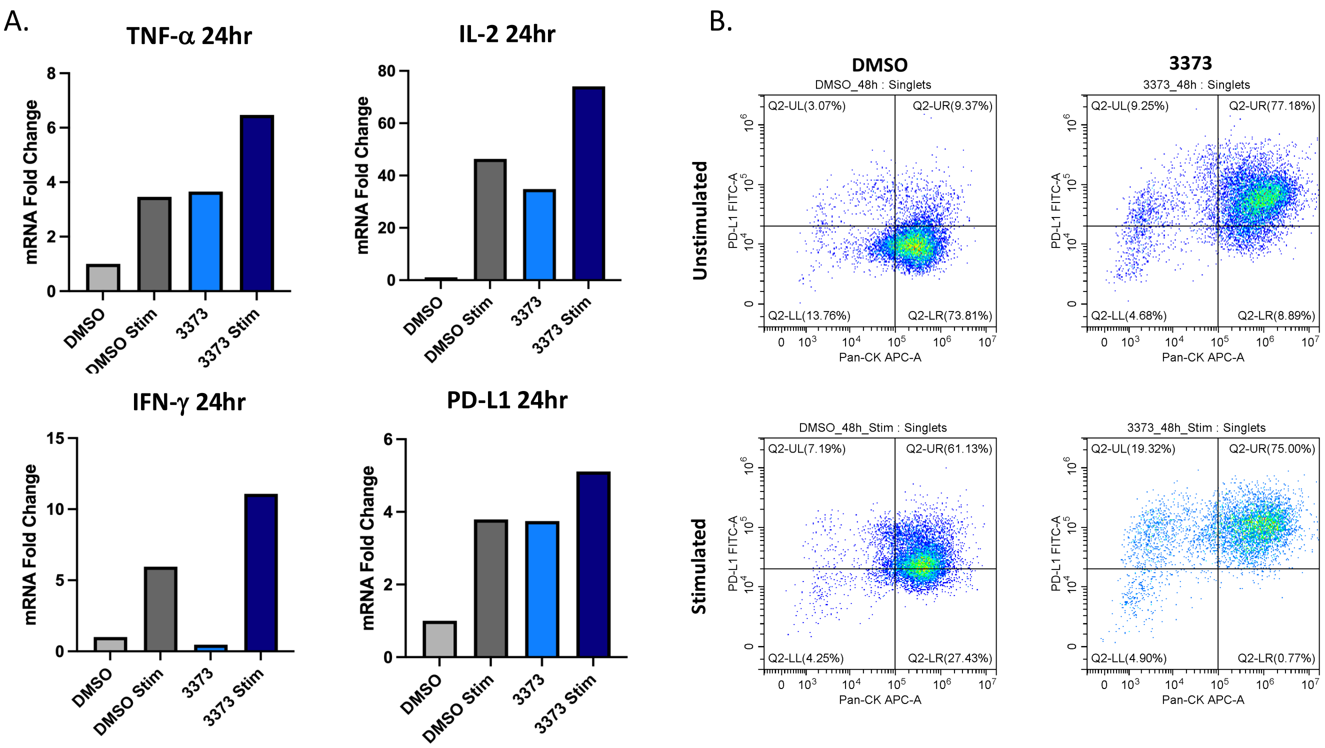


**Fig S3. NUC-3373 pre-treatment increases expression of cytokines and PD-L1 in PBMC/HCT116 cocultures:** HCT116 cells were pre-treated with either 10 μM NUC-3373 or DMSO for 24 hours before coculture with patient-derived PBMCs with or without stimulation with anti CD3/28. **A.** Representative gene expression of IL-2, TNF-α, IFN-γ, and PD-L1 in cocultures was determined by qPCR at 24hr. Gene expression plotted as fold change relative to 24 hours DMSO control. qPCR data was analysed using the ΔΔCt method. **B.** Flow cytometry data for PBMC/HCT116 cells stained for cell surface PD-L1 at 48 hours of coculture. Additional staining of intracellular pan-cytokeratin was used to differentiate HCT116 from PBMC.

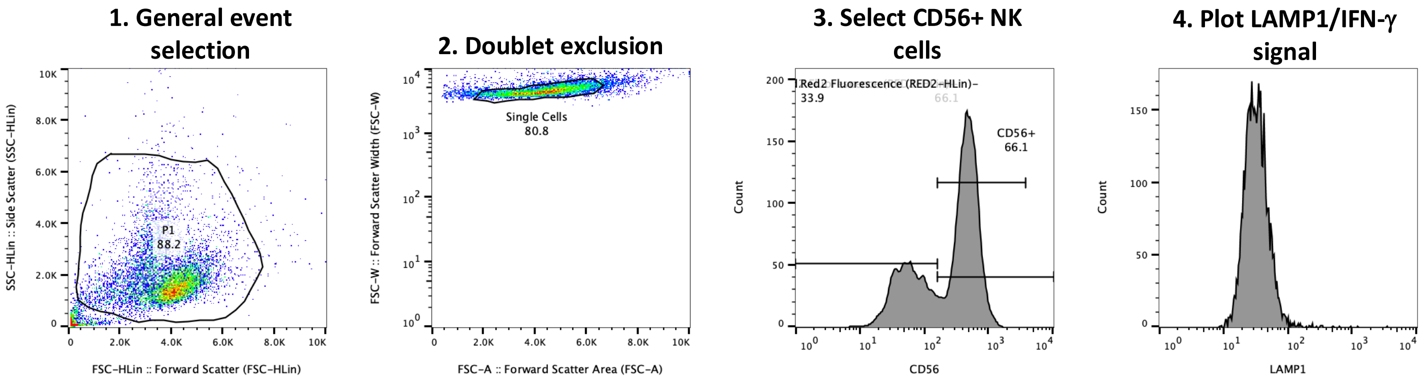


**Fig S4.** Flow cytometry gating strategy used for assessment of LAMP1 expression on CD56+ NK cells.

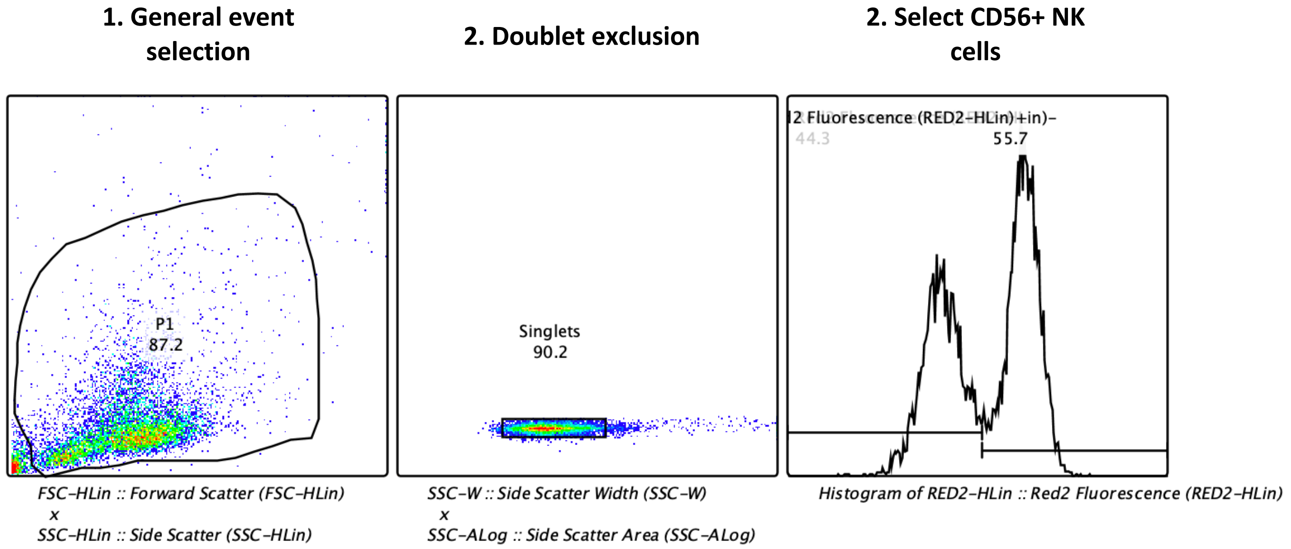


**Fig S5.** Flow cytometry gating strategy used for assessment of IFN-γ expression in CD56+ NK cells.

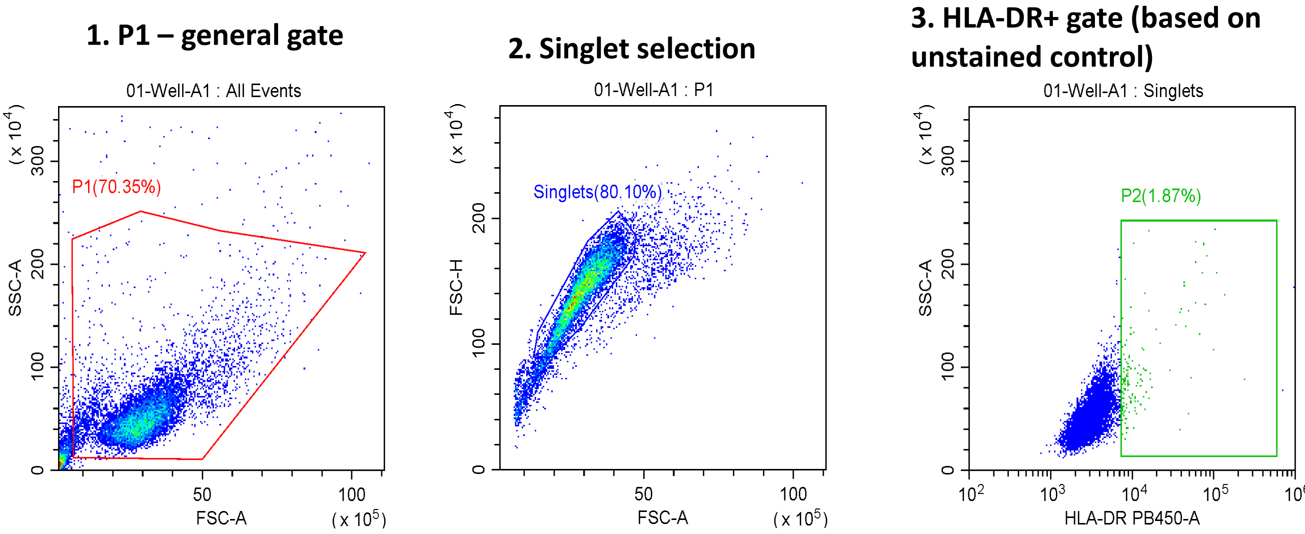


**Fig S6.** Flow cytometry gating strategy used for assessment of HLA-DR surface expression on CRC cells.

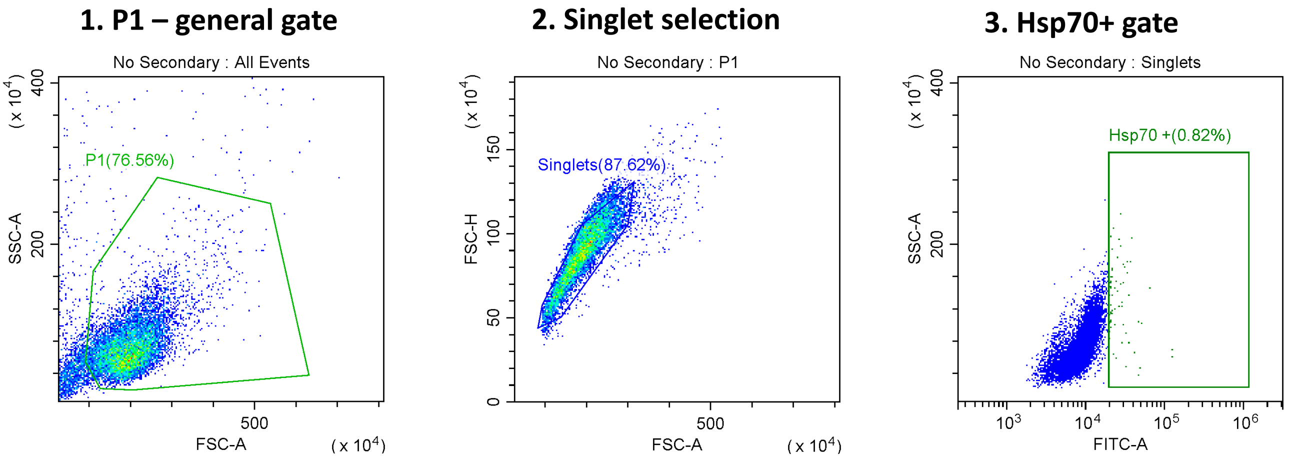


**Fig S7.** Flow cytometry gating strategy used for assessment of Hsp-70 surface expression on CRC cells.

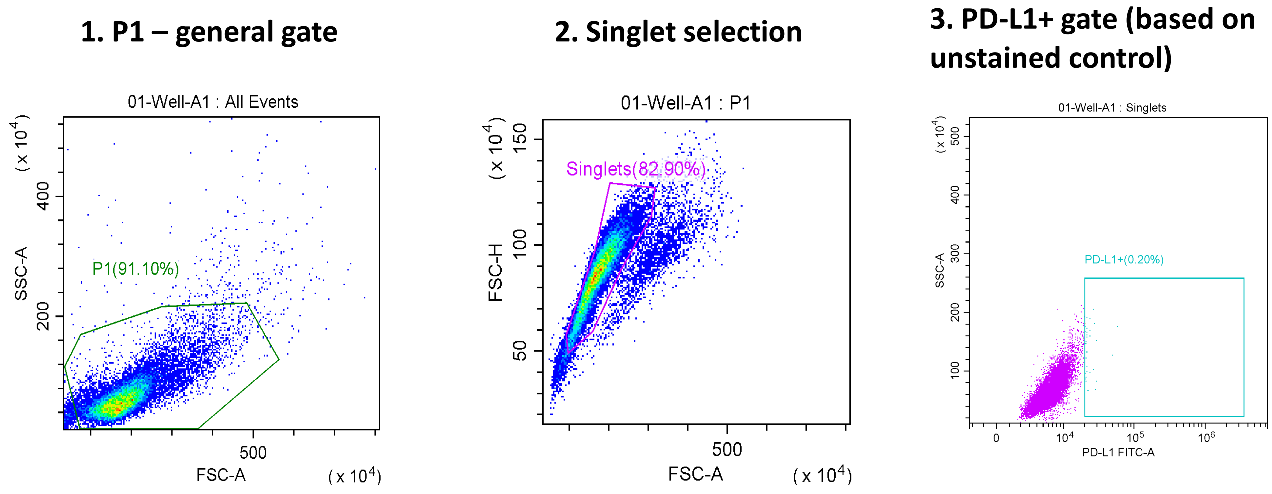


**Fig S8.** Flow cytometry gating strategy used for assessment of PD-L1 surface expression on CRC cells.
